# Dietary protein mediates terminal investment in egg quantity or quality following bacterial gut infection in Drosophila

**DOI:** 10.1101/489625

**Authors:** Ali L. Hudson, Joshua P. Moatt, Pedro F. Vale

**Affiliations:** Institute of Evolutionary Biology, School of Biological Sciences, University of Edinburgh Edinburgh EH9 3FL

**Keywords:** terminal investment, oral infection, dietary protein, fecundity compensation, *Drosophila melanogaster*, *Pseudomonas aeruginosa*

## Abstract

Organisms have evolved a range of behavioural and physiological responses which minimize the impact of infection on fitness. When future reproductive potential is threatened, for example, as a result of pathogenic infection, the terminal investment hypothesis predicts that individuals will respond by investing preferentially in current reproduction. Terminal investment involves reallocating resources to current reproductive effort, so it is likely to be influenced by the quantity and quality of resources acquired through diet. Dietary protein specifically has been shown to impact both immunity and reproductive output in a range of organisms, but its impact on terminal investment during infection is unclear. We tested the effect of dietary protein on terminal investment in the fruit fly *Drosophila melanogaster* following oral exposure to the opportunist bacterial pathogen *Pseudomonas aeruginosa*. Oral exposure to bacteria triggered an increase in reproductive investment, but we find that the nature of the terminal investment strategy depended on the level of dietary protein. Flies feeding on a high protein diet increased the number of eggs laid when exposed *to P. aeruginosa*, while flies fed an isocaloric, lower protein diet did not increase the number of eggs laid but instead showed an increase in egg-to-adult viability following infection. We discuss the importance of considering diet and natural routes of infection when measuring non-immunological defenses.

## Introduction

The life histories of all organisms are constrained by trade-offs, arising from the differential allocation of limited resources (Kirkwood, 1977; Stearns, 1992). For example, investing in current reproduction may be costly if it reduces the resources available for other somatic functions, such as growth, tissue repair or mounting an immune response (Schwenke *et al.*, 2016). The optimal resource allocation strategy will vary according to individual condition and environmental context, and a key trade-off is that between current and future reproduction (Williams, 1966; Holliday, 1989). When future reproductive potential is threatened, for example, as a result of pathogenic infection, reserving resources by spreading reproductive investment over multiple breeding attempts may result in reduced fitness relative to investing resources in current reproduction. The terminal investment hypothesis predicts that individuals will respond to such cues of impending sterility or mortality by increasing investment in current reproduction (Minchella & Loverde, 1981; Clutton-Brock, 1984; Thornhill *et al.*, 1986).

Terminal investment may take the form of increased early reproductive output, early maturation, or an increase in other forms of reproductive investment such as mating effort or parental care (Duffield *et al.*, 2017). Terminal investment has been observed in diverse animal and plant taxa in response to a wide range of cues (reviewed in Duffield *et al.*, 2017), including resource availability (Kim & Donohue, 2011), injury (Morrow *et al.*, 2003) non-pathogenic immune stimulation (Bonneaud *et al.*, 2004; Jacot *et al.*, 2004; Hanssen, 2006) and infection by lethal (Waldman *et al.*, 2016; Gupta *et al.*, 2017a), sub-lethal (Roznik *et al.*, 2015; Gupta *et al.*, 2017a), or sterilizing (Minchella & Loverde, 1981; Chadwick & Little, 2005; Vale & Little, 2012) pathogens. Because it increases host fitness during infection without directly reducing pathogen burdens, terminal investment acts to increase host disease tolerance, and has been described as an adaptive, non-immunological defense from infection (Parker *et al.*, 2011; Kutzer & Armitage, 2016a).

Terminal investment involves a reallocation of resources from other somatic functions to current reproductive effort, and thus is likely to be influenced by the quantity and quality of resources acquired through diet. Diet is known to affect both fecundity and immunity across a wide range of species (Lochmiller & Deerenberg, 2000; Field *et al.*, 2002; Lee *et al.*, 2008; Maklakov *et al.*, 2008; Jensen *et al.*, 2015; Schwenke *et al.*, 2016). Protein in particular is a key resource for growth, development and reproduction (Mirth *et al.*, 2019). Fruit flies *(Drosophila melanogaster)* produce more eggs on protein rich diets and these eggs are more likely to be viable (Drummond-Barbosa & Spradling, 2001; Lee *et al.*, 2008; Lihoreau *et al.*, 2016; Mirth *et al.*, 2019). Egg protein content is influenced directly by dietary protein (Kutzer & Armitage, 2016b; Mirth *et al.*, 2019) and has been shown to correlate with hatchling size (Stahlschmidt *et al.*, 2013). Egg protein content may additionally be subject to trade-offs against the immune response, as evidenced by immune challenged female mosquitoes *(Anopheles gambiae)* laying eggs with lower protein content (Ahmed *et al.*, 2002). Despite these findings, few studies have investigated how host diet or specific nutrients may influence the extent of terminal investment (Jacot *et al.*, 2004; Krams *et al.*, 2015).

In the present study we tested the effect of dietary protein on terminal investment in the fruit fly *D. melanogaster*. A previous study of systemic infection in *Drosophila* reared flies on either a standard or reduced protein diet but did not find any evidence for increased reproductive output following infection on either diet (Kutzer & Armitage, 2016c). Due to the expected tradeoff between reproduction and immunity, and the elevated protein requirements of oogenesis, we hypothesized that terminal investment would be more likely to occur on a high protein diet. We exposed female flies orally to the bacterial pathogen *Pseudomonas aeruginosa* in order to establish an enteric infection. We placed flies on a standard cornmeal-sugar-yeast Lewis diet (Lewis, 2014) or on a modified, isocaloric, high protein diet, and measured reproductive outputs that allowed us to assess the role of dietary protein on the reproductive quantity (the number of eggs laid) and also on the quality of those eggs (the number of eggs that eclosed as viable offspring).

## Methods

### Fly lines and rearing conditions

We used ten lines from the *Drosophila* Genetic Reference Panel (DGRP): RAL-59, RAL-75, RAL-138, RAL-373, RAL-379, RAL-380, RAL-502, RAL-738, RAL-765 and RAL-818 (Mackay *et al.*, 2012). All lines were previously cleared of *Wolbachia* infection, which is known to confer protection against enteric bacterial infection by *P. aeruginosa* (Gupta *et al.*, 2017b). Prior to the experiment, all lines were housed in plastic vials (*Φ* 25mm, height 95mm) plugged with non-absorbent cotton wool on a standard undyed Lewis diet (Lewis, 2014) and maintained under identical conditions of 12:12 light:dark regimes at 25°C for minimum 3 generations. Stocks were kept at 10-20 adult flies per vial and allowed to lay for 24 hours before being removed. Flies laid for the experimental generations were density controlled by adding 15 female and 2 male flies to each vial for 24 hours. Eggs laid during this period were allowed to develop for 14 days at 25°C. The resulting adults were lightly sedated with CO_2_, 14 days after the parents had been introduced to lay eggs. Two density-controlled vials were set up for each line by placing 15 females and 2 males on standard Lewis medium, where they were kept for 24 hours (±2 hours) to ensure maturity and mating had occurred prior to the experiment.

### Diet treatments and experimental setup

Two diets of differing protein levels were used (Table S1). A standard Lewis diet of roughly 14% protein was chosen, as this is frequently employed in laboratory experiments involving *Drosophila*. The second diet was a Lewis diet modified to contain approximately double the amount (~31 %) protein, as it was shown to induce significantly higher egg laying in *Drosophila* (Lee *et al.*, 2008; Jensen *et al.*, 2015). Protein quantity was manipulated by increasing the yeast component, while carbohydrate was reduced by decreasing the sugar to maintain an approximately isocaloric diet. Both diets were dyed with Brilliant Blue FCF E133 (SIgma) to increase contrast between the eggs and the food during egg counts. The experiment used a 2x2x10 fully cross-factored design, with two levels of infection status (infected and uninfected), two diets (normal 7% and high 14%), and ten fly lines. Ten, individually housed, replicate flies from each line were subject to each treatment. This resulted in a total of 400 flies, 40 per line, being used, with 200 flies for each level of the factors diet and infection, and 100 flies for each diet-infection status combination. The experiment was split evenly over two blocks, with 5 replicates per line per treatment group in each block.

### *Pseudomonas aeruginosa* culture and oral infection protocol

*P. aeruginosa* reference strain PA14 is a gram-negative bacterium known to cause mortality in a range of species, including *D*. melanogaster (Apidianakis & Rahme, 2009; De Soyza *et al.*, 2013). Bacterial growth for fly oral infection was carried out as described in previously (Siva-Jothy *et al.*, 2018). Briefly, a 200μl stock culture of PA14 (optical density at 600nm, OD_600_=1) frozen at -80°C in 25% glycerol was introduced to a 50ml falcon tube containing 20ml of sterile LB broth (Fisher Scientific BP1426), and shaken overnight at 140rpm and 30°C. To produce the large volume and high concentration of bacteria needed for infection, subcultures were taken from the overnight cultures by introducing 3ml of culture into 297ml of sterile LB broth. These were shaken at 140rpm for 7-8 hours at 37°C, and monitored until they reached OD_600_=0.6 to 0.8, indicative of the exponential growth phase. Each subculture was divided into 50ml falcon tubes, containing 30ml of subculture each and centrifuged at 2,500xg at 4°C for 20 minutes to precipitate the bacteria. The majority of the supernatant was discarded, except for the final ~2ml, in which the pellet was resuspended by vortexing at a high speed for 2-3 minutes. These suspensions were transferred to a single falcon tube which was centrifuged again as above, and the supernatant discarded. The pellet from each subculture was resuspended in 5% sucrose solution to achieve an OD_600_=25.

For oral infection, flies were starved for 7-8 hours prior to infection by tipping into foodless vials, bunged with absorbent cotton wool moistened with distilled water to prevent dehydration. In the 24 hours preceding the infection protocol, 500μl of sugar agar (20g of agar powder and 84g of brown sugar, dissolved in 1l distilled H_2_O and heated) was added to the lid of a 7ml Bijou tube (Fisher Scientific 129A). Once firm, a 20mm filter paper disc was placed on the agar, and the bijous were sealed for overnight storage at 4°C, and returned to room temperature before use. Immediately before infection, 80μl of the PA14 suspension (OD_600_=25 as described above), or 5% sucrose for the control, was pipetted onto the filter disc and allowed to dry for 20 minutes. The starved flies were lightly sedated with CO_2_, transferred individually into a bijou and kept overnight (~16 hours) at 25°C to ingest bacteria. They were then tipped onto their designated diets. To confirm infection and the absence of contamination of the control flies, we prepared 20 additional flies from each line and quantified bacterial growth at the end of the infection period. These flies were placed individually into 1.5ml Eppendorf tubes and surface sterilized for 30-60 seconds in 100μl 70% ethanol, then washed twice in 200μl of distilled water. 5μl of the second wash was plated on an LB agar plate (Fisher Scientific BP1426) and another 5 μl on *Pseudomonas* Isolation Medium (PIM) (Sigma-Aldrich P2102) plate, to confirm surface sterilization. The flies were then placed in 1.0ml of Phosphate Buffer Solution (PBS) and microcentrifuged at 5000rpm for 1 minute before the PBS was removed and the fly homogenized in 200μl of LB broth using a micropestle and handheld motor. This homogenate was plated on sterile PIM agar, and incubated at 29°C for 2-3 days to confirm infection. This confirmed infection in all PA14 treated flies tested, and found no evidence of contamination of control flies.

### Fecundity and survival following infection

Following infection, the flies were housed individually on either the standard Lewis diet, or the modified higher protein diet (described above), and maintained at 25°C on a 12:12 light:dark cycle. All flies were tipped onto fresh food of the same diet every day for seven days, when their survival was recorded, and the number of eggs laid counted under a microscope. Survival was recorded for an additional 3 days after egg counts concluded. To assess egg-to-adult viability, eggs laid on days 1-3, 5 and 7 were incubated for 16 days at 25°C, and the number of eclosed offspring were counted.

### Analysis

Analysis and plots were performed using R version 3.4.3 (Core Team, 2017) using the packages *lme4* (Bates *et al.*, 2015) and *survival* (Therneau, 2015). All models include the random effect of individual nested within line to account for repeated measures across individuals and lines. Daily egg production and number of eclosed offspring were analyzed via generalized linear mixed effects models (GLME). Models fitted diet, infection status, day and their interactions, alongside block as categorical fixed effects. To control for overdispersion within the data, row ID was included as a random effect in both models. Egg-to-adult viability was analysed using a binomial GLME, with the number of eggs that eclosed and the number which did not eclose bound and treated as the response variable, i.e. the proportion of eggs eclosing. Diet, infection status, day and their interactions were treated as fixed effects as well as block. To account for potential density effects, the total number of eggs present in the vial was included as a random effect. To understand any life-history changes induced solely by diet, infection and its interactions were dropped and all models were rerun on control flies only. Full R code for all analysis is available in *<u>supplementary materials /DRYAD</u>*.

## Results

### Life-history changes due to dietary protein in uninfected flies

Before examining the effects of dietary protein on terminal investment, it is important to assess its effects on reproductive output in healthy flies. As anticipated, flies reared on the high protein diet produced more eggs than those on the standard diet (Figure 1), and these eggs showed higher viability (Figure 2), resulting in more eclosed adult offspring per fly each day under high protein compared to on the standard diet (Figure 3; Table 1). The number of eggs laid each day increased over the course of the experiment when flies were reared on the high protein diet, but this increase was not as evident under the standard protein diet (Figure 1, light blue bars; Table 1, ‘Diet x Day’ interaction). Diet-dependent temporal dynamics were also evident for the number of viable offspring (Figure 2; Table 1, ‘Diet x Day’ interaction). We found that the genetic background of flies contributed significantly to the variance in both the number of eggs laid and in the proportion of these eggs that resulted in the eclosion of viable offspring (Table 1 “line” effect; Figures S1–S3).

**Figure 1.**
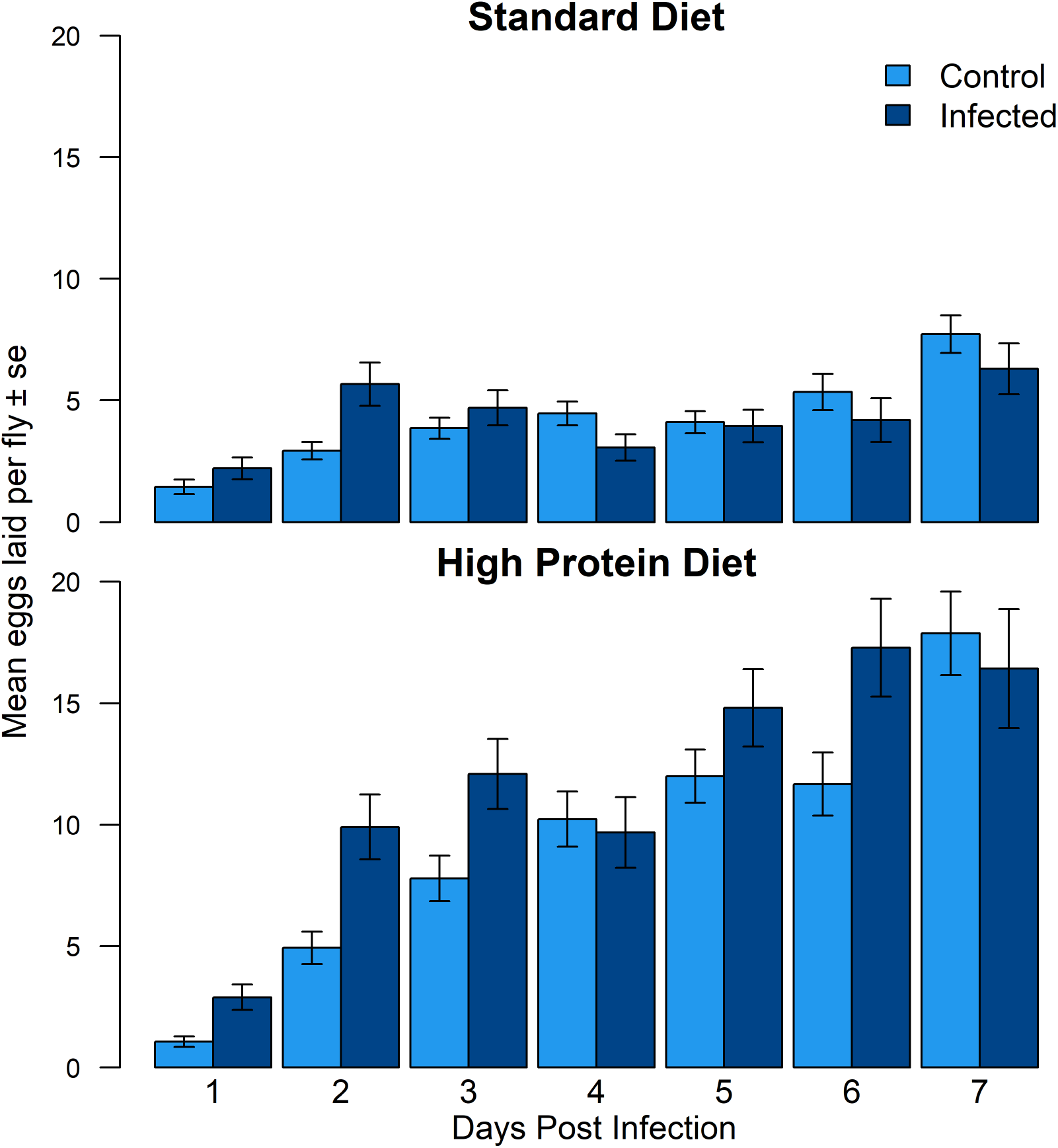
Egg Production. Mean number of eggs laid per fly by control flies (light blue) and infected flies (dark blue) on the first seven days following infection on the standard Lewis diet (top) and the modified high protein diet (bottom).

**Figure 2.**
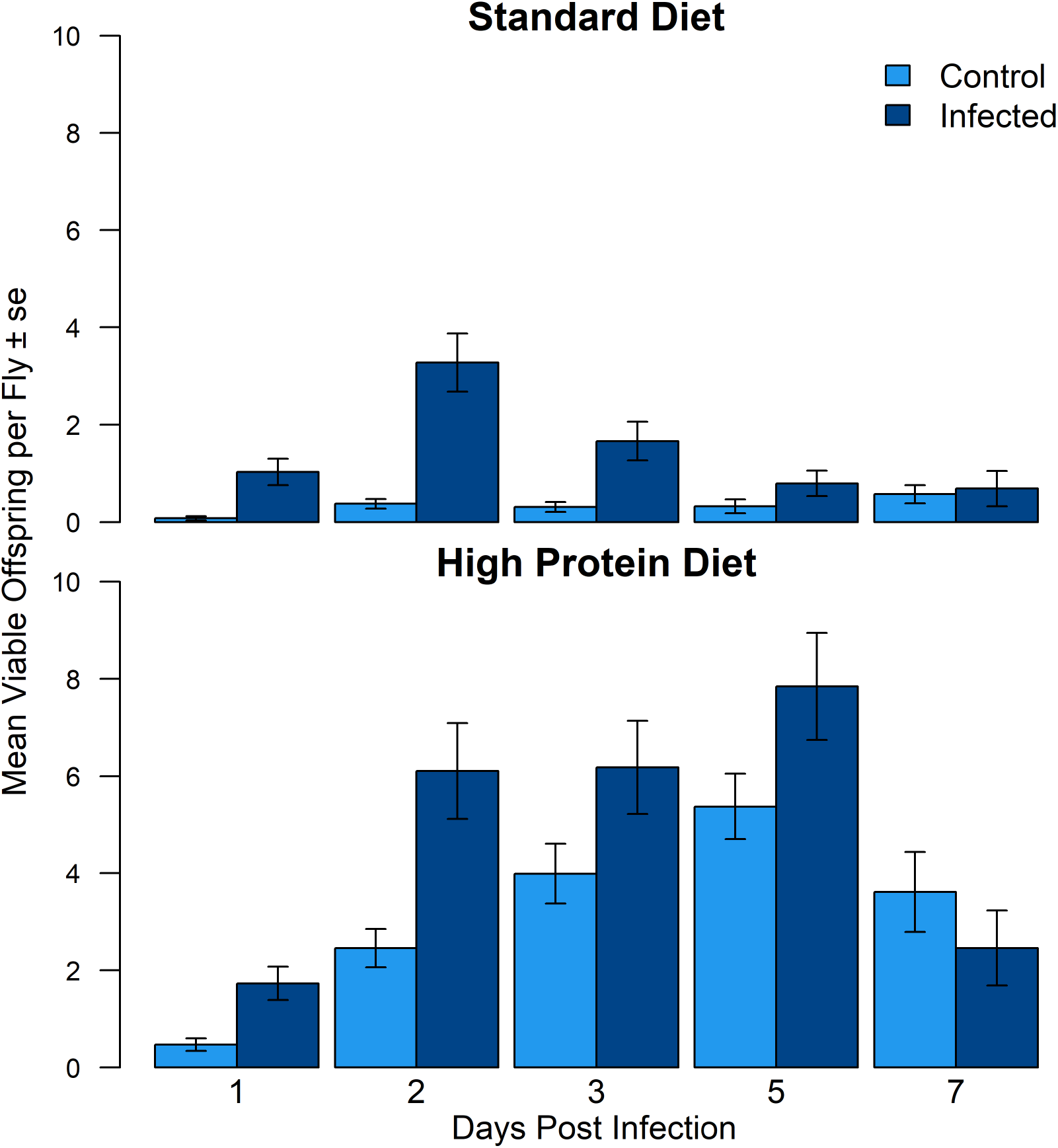
Total Viable Offspring. Mean number of eclosed offspring per fly by control flies (light blue) and infected flies (dark blue) over seven days following infection on the standard Lewis diet (top) and the modified high protein diet (bottom).

**Figure 3.**
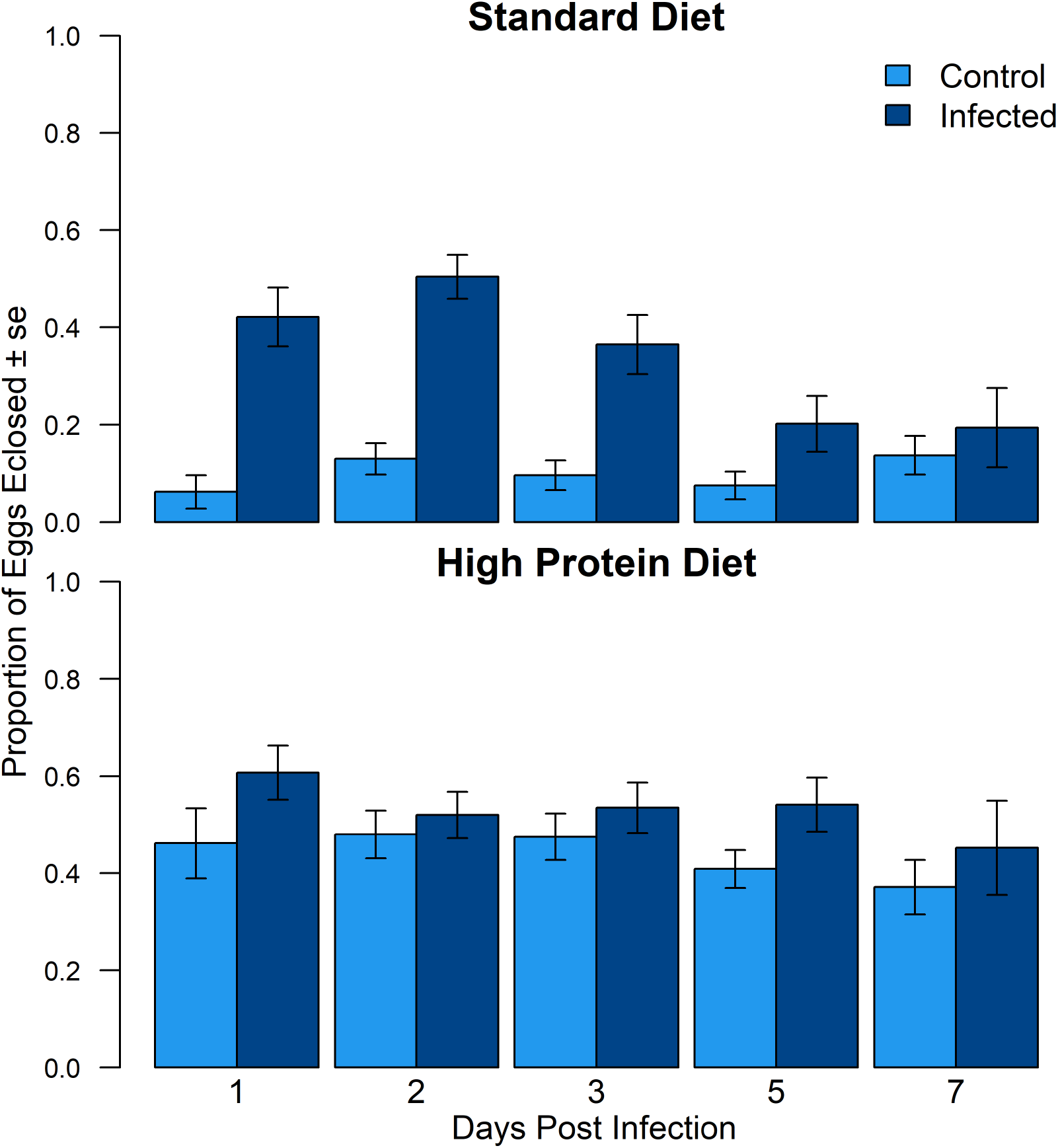
Egg-Adult Viability. Proportion of eggs which eclosed laid by control flies (light blue) and infected flies (dark blue) over seven days following infection on the standard Lewis diet (top) and the modified high protein diet (bottom).

**Table 1:**
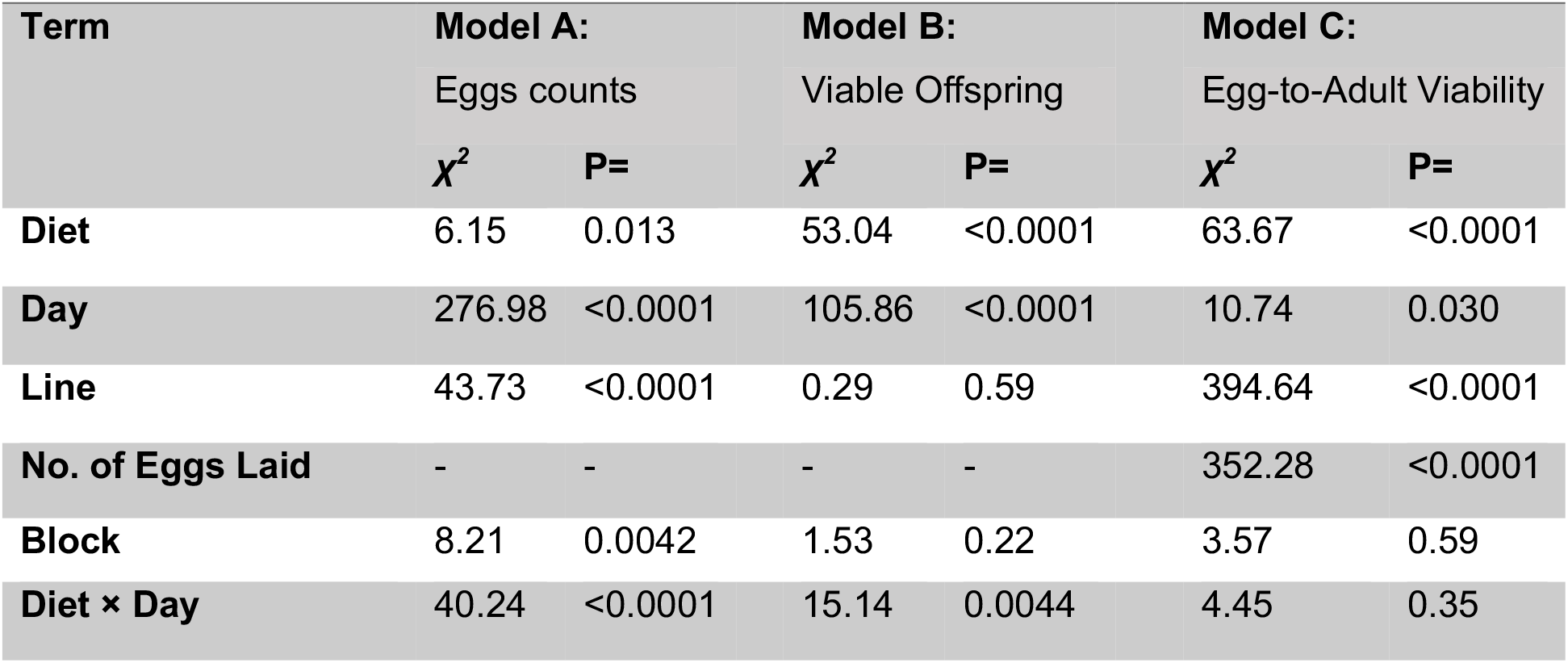
Summarized models of control non-infected flies only.

| Term | Model A: |  | Model B: |  | Model C: |  |
| --- | --- | --- | --- | --- | --- | --- |
|  | Eggs counts |  | Viable Offspring |  | Egg-to-Adult Viability |  |
| | $\chi^2$ | P= | $\chi^2$ | P= | $\chi^2$ | P= |
| **Diet** | 6.15 | 0.013 | 53.04 | <0.0001 | 63.67 | <0.0001 |
| **Day** | 276.98 | <0.0001 | 105.86 | <0.0001 | 10.74 | 0.030 |
| **Line** | 43.73 | <0.0001 | 0.29 | 0.59 | 394.64 | <0.0001 |
| **No. of Eggs Laid** | - | - | - | - | 352.28 | <0.0001 |
| **Block** | 8.21 | 0.0042 | 1.53 | 0.22 | 3.57 | 0.59 |
| **Diet × Day** | 40.24 | <0.0001 | 15.14 | 0.0044 | 4.45 | 0.35 |

### Increased oviposition in infected flies on high protein diet

Flies exposed orally to *Pseudomonas aeruginosa* experienced significantly higher mortality than control flies and the rate of mortality did not differ with diet (Figure S4). Most mortality (approximately 40%) occurred within 1-3 days following oral exposure, reaching 50% by the end of the experiment. The genetic background of the flies explained a significant proportion of variance in the number of eggs laid (Table 2 “line” effect; Figures S1). Flies that were exposed to *P.aeruginosa* laid significantly more eggs than those exposed to a control solution, but only when fed the high protein diet (Figure 1; Table 2, Model 1 ‘Diet x Infection Status’). Averaged over all days, exposed flies on the high protein diet laid 9.3 eggs per day, compared to 7.6 laid per day by control flies on the same diet.

**Table 2:** Summarized models for infected and control flies.

| Term | Model 1:<br>Eggs counts |  | Model 2:<br>Viable Offspring |  | Model 3:<br>Egg-to-Adult Viability |  |
| --- | --- | --- | --- | --- | --- | --- |
| | $\chi^2$ | P= | $\chi^2$ | P= | $\chi^2$ | P= |
| **Diet** | 17.26 | <0.0001 | 69.87 | <0.0001 | 73.55 | <0.0001 |
| **Infection** | 0.0223 | 0.88 | 30.41 | <0.0001 | 40 | <0.0001 |
| **Day** | 307.14 | <0.0001 | 123.6 | <0.0001 | 61.69 | <0.0001 |
| **Line** | 71.7 | <0.0001 | 21.62 | <0.0001 | 98.92 | <0.0001 |
| **No. of Eggs Laid** | - | - | - | - | 29.79 | <0.0001 |
| **Block** | 12.35 | <0.001 | 1.71 | 0.19 | 13.06 | <0.001 |
| **Diet × Infection** | 4.45 | 0.035 | 6.54 | 0.011 | 24.99 | <0.0001 |
| **Diet × Day** | 76.37 | <0.0001 | 43.22 | <0.0001 | 65.52 | <0.0001 |
| **Infection × Day** | 75.23 | <0.0001 | 37.12 | <0.0001 | 23.12 | <0.001 |
| **Diet x Infection × Day** | 6.88 | 0.33 | 1.39 | 0.85 | 22.37 | <0.001 |

### Egg viability is increased in infected flies, regardless of diet

While increasing the number of eggs following exposure to a pathogen is a clear indication of terminal investment, more eggs will only translate into increased fitness if they are capable of developing into viable adult offspring. As expected, infected flies on the high protein diet produced a greater number of viable offspring than those on the standard diet (Figure 3; Table 2, Model 2, ‘Diet x Infection’), reflecting their higher egg laying. However, the higher number of viable adult offspring from infected flies was not only a result of increased egg laying, but also due to an increase in egg-to-adult viability, which was higher in infected flies relative to control flies. Flies on the standard diet showed a larger increase in viability following infection than those on the high protein diet, peaking 2 days post-infection. Both the total number of eclosed offspring and the egg-to-adult viability differed between fly lines (Table 1 “line” effect; Figures S2–S3).

## Discussion

Organisms have evolved an array of strategies to minimize the impact of infection on fitness, including behavioral avoidance of infection (Curtis, 2014; Vale *et al.*, 2018), and mechanisms that either mediate pathogen clearance or that minimize the damage caused by pathogen exploitation (Gupta & Vale, 2017; Soares *et al.*, 2017; Lissner & Schneider, 2018). These defense mechanisms are likely to be costly to maintain and deploy (Moret & Schmid-Hempel, 2000; Armitage *et al.*, 2003; Bonneaud *et al.*, 2003; Labbé *et al.*, 2010; Duncan *et al.*, 2011; Auld *et al.*, 2013; Susi & Laine, 2015; Vale *et al.*, 2015), and therefore rely heavily on the acquisition of dietary nutrients, their transformation into energy resources, and the appropriate allocation of these resources to different life-history traits (Schwenke *et al.*, 2016).

We investigated the effect of dietary protein on terminal investment in response to infection, a form of non-immunological defense that mitigates the potential fitness losses of infection by increasing reproductive investment (Parker *et al.*, 2011; Kutzer & Armitage, 2016a). We found that oral infection by *P. aeruginosa* was sufficient to trigger a shift in reproductive investment, recapitulating similar increases in reproductive output in *D. melanogaster* following sub-lethal viral infections (Gupta *et al.*, 2017a). However, here we observed that the nature of the terminal investment strategy depended on the availability of dietary protein. Flies feeding on a high protein diet invested terminally in the quantity of eggs, while flies fed a lower protein diet increased investment in the quality (viability) but not quantity of their eggs.

While there is a considerable amount of work showing that protein levels affect reproductive output and immunity (reviewed in Schwenke *et al.*, 2016), the role of diet on the ability to terminally invest following exposure to pathogens has received less attention. In one study, diet-restricted male mealworm beetles (*Tenebrio molitor*) were found to invest terminally in attractive sex odours at the expense of a resistant encapsulation response to a nylon implant (Krams *et al.*, 2015a). In other work, reduced investment in mate calling by male crickets injected with bacterial lipopolysaccharides was alleviated by food supplementation (Jacot *et al.*, 2004), suggesting that reproductive investment following infection can be augmented by dietary supplementation.

In Drosophila, both the quantity of protein per egg and the quantity of eggs produced are influenced by dietary protein availability (Mirth *et al.*, 2019). Female *D. melanogaster* typically weigh 800-1100μg (wet weight, Jumbo-Lucioni et al., 2010), and lay eggs containing approximately 10-12μg of protein each (Kutzer and Armitage, 2016). The highest laying fly in this study produced 172 eggs over 7 days, representing about 2000μg of protein invested in egg production, or ~200 % of the fly’s wet weight, which underlines the importance of dietary protein for oogenesis. In the current experiment, protein was clearly the factor limiting investment in increased egg production, since in line with the results of other studies (Drummond-Barbosa & Spradling, 2001; Lee *et al.*, 2008; Kutzer & Armitage, 2016c), flies on the elevated protein diet produced more eggs than flies on the standard medium. The lack of terminal investment in the number of eggs under the lower level of protein may therefore be a result of the necessary protein being unavailable on the standard diet. It is therefore plausible that other studies where terminal investment has not been observed were a result of insufficient protein being available to terminally invest in increased reproduction (e.g Kutzer & Armitage, 2016b).

Investing in increased egg production is one way organisms can improve their number of surviving offspring, but another is to ensure that the offspring produced are viable. We took egg-to-adult viability to reflect egg quality, counting both the number of eggs laid by a fly on a given day, and the number of those eggs which eclosed to adults within 16 days. The greatest increase in egg-to-adult viability following infection was observed in eggs laid by flies on the standard diet, whereas those laid by infected flies on the standard diet were more numerous but not more viable than those of uninfected controls. Previous work has found that flies raised on a poor diet produce heavier eggs, and produce offspring that themselves are more resistant to poor nutrition than those of flies raised on a standard diet (Vijendravarma *et al.*, 2010). This suggests that flies may be subject to a protein allocation trade-off between per-egg protein allocation, and number of eggs produced, and that payoffs of this trade-off vary according to the quality of food available. In a situation of low protein availability, it may be better to invest what little protein is available in a smaller number of eggs to improve offspring viability.

The precise mechanisms by which changes in diet affect reproductive traits following infection are difficult to disentangle. Dietary protein provides both the raw material for egg production, as well as influencing complex signalling pathways which determine investment in egg production (Mirth, Alvez & Piper, 2019) Our results showed that flies on the standard diet could produce eggs with higher viability but did not invest in doing so in the absence of infection. This suggests that raw materials were available to produce more viable eggs, but signalling pathways controlling investment in egg quality were influenced by limited protein availability to reduce this investment. Recent research has highlighted the roles played by juvenile hormone and ecdysone levels as well as insulin signalling in regulating egg production in response to nutritional states (Mirth et al. 2019). Additionally, bacterial derived peptidoglycans have been shown to activate NF-kB signalling pathways in octopaminergic neurons, resulting in changes in egg laying (Kurz *et al.*, 2017). Interactions between these pathways signalling nutritional and infection status may therefore underlie protein-mediated plasticity in terminal investment. Future work should investigate these interactions and attempt to characterise their potential as a mechanism by which organisms can pursue optimal strategies under differing nutrient availabilities.

Compared to previous work on terminal investment, particularly in insect systems, a unique aspect of this study was the infection method. We chose to establish a gut infection because we were investigating an evolved adaptive response to infection, and oral infection by Pseudomonas is believed to be more common in the wild than infection via septic route employed in many other studies (Jacot *et al.*, 2004; Reaney & Knell, 2010; Duffield *et al.*, 2017). Other work has shown that the evolutionary response of *D. melanogaster* to *Pseudomonas* infection is specific to the route of infection (Martins *et al.*, 2013), and that antibacterial protection by *Wolbachia* occurs during oral but not systemic infections (Gupta *et al.*, 2017b). These results suggest that selection to cope with oral Pseudomonas infection has been stronger, which may explain why previous works which often employed systemic infections have not detected a similar terminal investment response (Kutzer & Armitage, 2016c).

In summary, we find that dietary protein can mediate the terminal investment strategy of flies following infection. This result places our current understanding of non-immunological defence from infection in an important ecological context, as environments where protein availability is variable may select for multiple resource-dependent strategies for limiting the impact of infection. Further research into the wider consequences of this plasticity on the population ecology of host species during infection, and the underlying physiological mechanisms of these responses is now needed. Combined, this will result in a clearer understanding of the broader ecological and evolutionary implications of fluctuating resource availability in natural populations.

## Acknowledgements

We are grateful to Katy Monteith and Jay-Russell Dennis for technical assistance, Angela Reid and Lucinda Rowe for preparation of all fly media and Eevi Savola for sharing information on diet composition. We also thank Jenny Regan, Darren Obbard, Craig Walling and respective lab members for constructive suggestions. P. Vale is grateful to funding and support from a Chancellor’s fellowship (University of Edinburgh) and a Branco Weiss fellowship from Society in Science.

## Supplementary Tables and Figures

**Table S1:** A comparison of the both diets with ingredients for approximately 11 of food, or enough for ~100 vials.

|  | Protein % | P:C ratio | Yeast (g) | Sugar (g) | Maize (g) | Agar (g) | Nipagin (ml) | Food Dye (g) | dH<sub>2</sub>O (l) |
| --- | --- | --- | --- | --- | --- | --- | --- | --- | --- |
| **Standard Diet** | 14% | 1:6 | 18.75 | 93.75 | 69.17 | 6.87 | 15 | 0.5 | 1 |
| **High Protein** | 31% | 1:2 | 49.45 | 63.05 | 69.17 | 6.87 | 15 | 0.5 | 1 |

**Figure S1.**
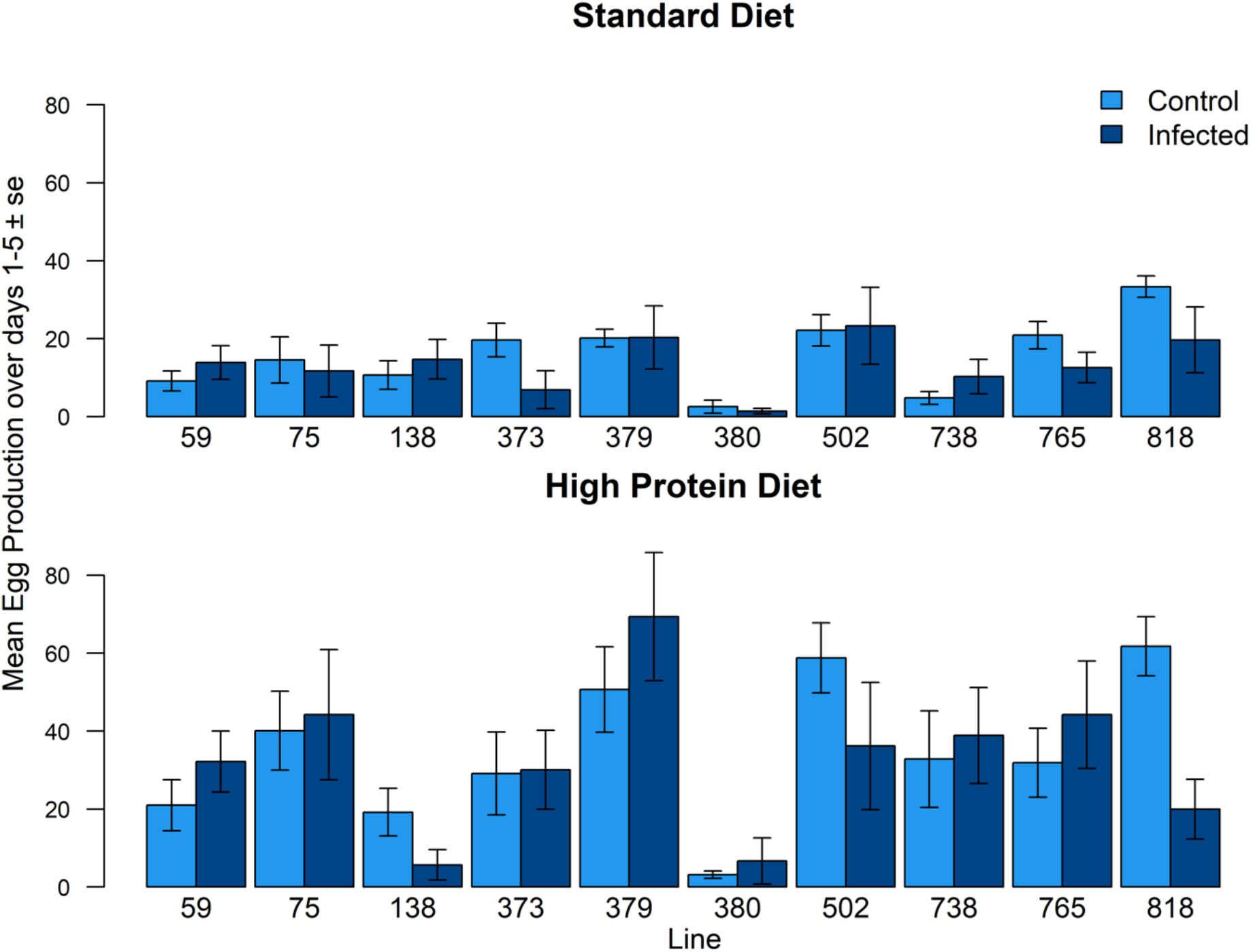
Mean daily egg production by line, diet, and infection status. Mean daily egg production per fly by control flies (light blue) and infected flies (dark blue) by line over the first five days following infection on the standard Lewis diet (above) and the modified high protein diet (below).

**Figure S2.**
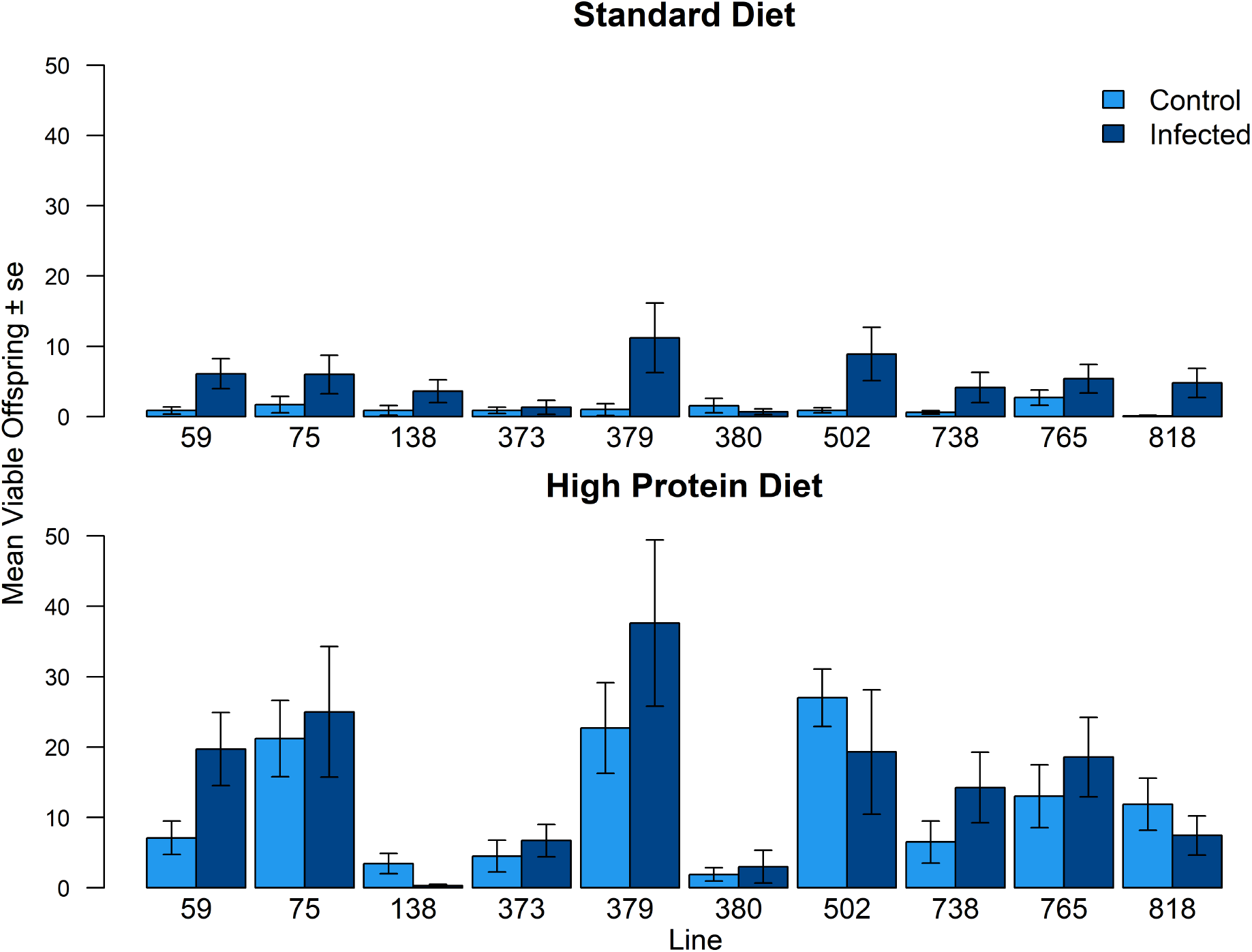
Mean viable offspring per fly per day, by line, diet, and infection status. Mean number of eclosed offspring produced over the first five days following infection per fly by control flies (light blue) and infected flies (dark blue) by line for flies on the standard Lewis diet (above) and the modified high protein diet (below).

**Figure S3.**
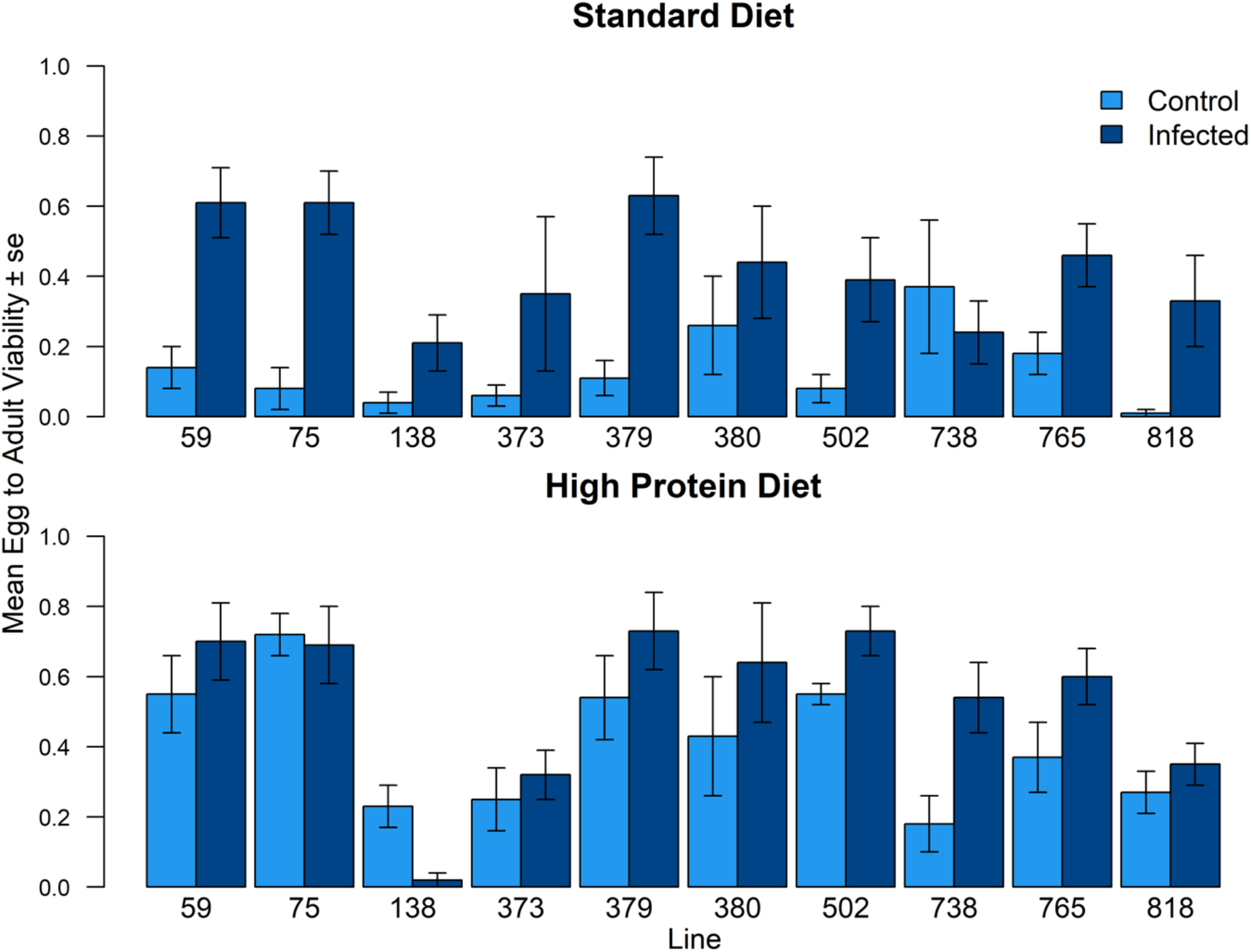
Mean egg-to-adult viability by line, diet, and infection status. Proportion of eggs laid which eclosed laid by control flies (light blue) and infected flies (dark blue) by line over the first five days following infection on the standard Lewis diet (above) and the modified high protein diet (below).

**Figure S4.**
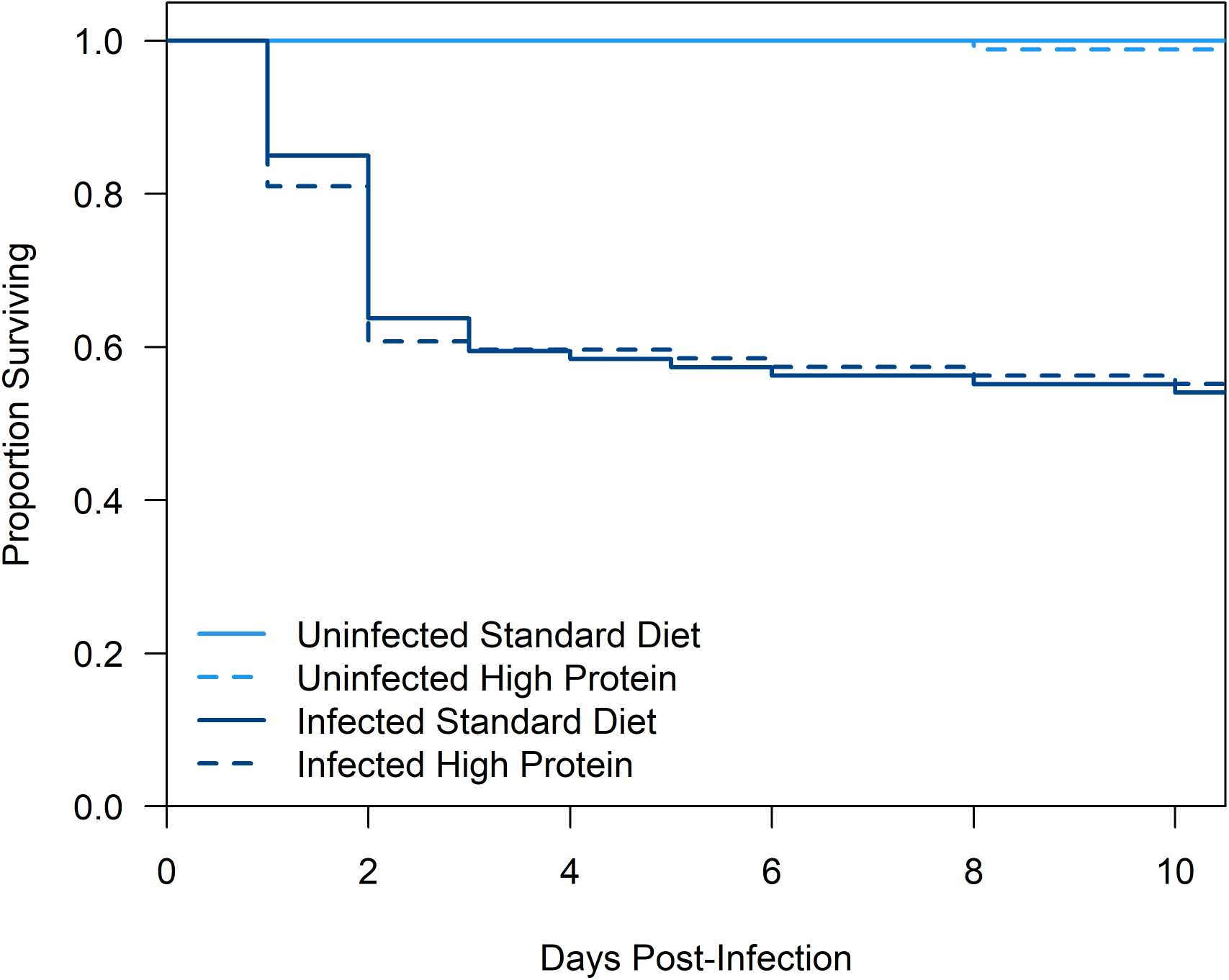
Kaplan Meier Plot of survival. Kaplan Meier plot of survival for control (light blue) and infected (dark blue) flies on the standard (solid line) and modified high protein (dashed line) diets over the first ten days following infection

## Supplementary Estimates SEs tables

| **Egg Production** | **Estimate (s.e.)** | **X<sup>2</sup>** | **P** |
| --- | --- | --- | --- |
| (Intercept) | -1.337 (0.339) |  |  |
| DietNormal | 0.417 (0.292) | 17.26 | < 0.001 |
| InfectionInfected | 0.874 (0.282) | 0.02 | 0.880 |
| Day2 | 1.516 (0.200) |  |  |
| Day3 | 2.030 (0.197) |  |  |
| Day4 | 2.368 (0.195) |  |  |
| Day5 | 2.777 (0.194) |  |  |
| Day6 | 2.548 (0.226) |  |  |
| Day7 | 2.063 (0.229) | 307.14 | < 0.001 |
| BlockB | 0.538 (0.152) | 12.35 | < 0.001 |
| DietNormal:InfectionInfected | -0.737 (0.401) | 4.45 | 0.035 |
| DietNormal:Day2 | -0.612 (0.281) |  |  |
| DietNormal:Day3 | -0.853 (0.278) |  |  |
| DietNormal:Day4 | -0.995 (0.276) |  |  |
| DietNormal:Day5 | -1.511 (0.276) |  |  |
| DietNormal:Day6 | -1.054 (0.322) |  |  |
| DietNormal:Day7 | -0.421 (0.324) | 76.37 | < 0.001 |
| InfectionInfected:Day2 | -0.156 (0.274) |  |  |
| InfectionInfected:Day3 | -0.367 (0.280) |  |  |
| InfectionInfected:Day4 | -1.183 (0.284) |  |  |
| InfectionInfected:Day5 | -0.851 (0.278) |  |  |
| InfectionInfected:Day6 | -0.723 (0.333) |  |  |
| InfectionInfected:Day7 | -2.027 (0.362) | 75.23 | < 0.001 |
| DietNormal:InfectionInfected:Day2 | 0.281 (0.391) |  |  |
| DietNormal:InfectionInfected:Day3 | -0.067 (0.404) |  |  |
| DietNormal:InfectionInfected:Day4 | 0.063 (0.413) |  |  |
| DietNormal:InfectionInfected:Day5 | 0.077 (0.408) |  |  |
| DietNormal:InfectionInfected:Day6 | -0.371 (0.492) |  |  |
| DietNormal:InfectionInfected:Day7 | 0.903 (0.511) | 6.88 | 0.330 |

| **Number of Viable Offspring** | **Estimate (s.e.)** | **X<sup>2</sup>** | **P** |
| --- | --- | --- | --- |
| (Intercept) | -2.435 (0.369) |  |  |
| DietNormal | -1.758 (0.558) | 53.04 | <0.0001 |
| InfectionInfected | 1.377 (0.368) | 30.41 | <0.0001 |
| Day2 | 1.778 (0.287) |  |  |
| Day3 | 2.260 (0.283) |  |  |
| Day5 | 2.760 (0.280) |  |  |
| Day7 | 2.002 (0.334) | 123.60 | <0.0001 |
| BlockB | 0.253 (0.194) | 1.71 | 0.190 |
| DietNormal:InfectionInfected | 1.092 (0.656) | 6.54 | 0.011 |
| DietNormal:Day2 | -0.090 (0.587) |  |  |
| DietNormal:Day3 | -0.886 (0.600) |  |  |
| DietNormal:Day5 | -1.534 (0.611) |  |  |
| DietNormal:Day7 | -0.056 (0.660) | 43.22 | <0.0001 |
| InfectionInfected:Day2 | -0.472 (0.376) |  |  |
| InfectionInfected:Day3 | -0.862 (0.386) |  |  |
| InfectionInfected:Day5 | -1.109 (0.384) |  |  |
| InfectionInfected:Day7 | -2.089 (0.507) | 37.12 | <0.0001 |
| DietNormal:InfectionInfected:Day 2 | 0.153 (0.689) |  |  |
| DietNormal:InfectionInfected:Day 3 | -0.005 (0.722) |  |  |
| DietNormal:InfectionInfected:Day 5 | -0.458 (0.749) |  |  |
| DietNormal:InfectionInfected:Day 7 | -0.499 (0.894) | 1.39 | 0.850 |

| **Egg-Adult Viability** | **Estimate (s.e.)** | **X2** | **P** |
| --- | --- | --- | --- |
| (Intercept) | 0.116 (0.389) |  |  |
| InfectionInfected | 0.603 (0.350) | 40.00 | <0.0001 |
| Day2 | -0.238 (0.281) |  |  |
| Day3 | -0.041 (0.273) |  |  |
| Day5 | -0.350 (0.276) |  |  |
| Day7 | -0.329 (0.300) | 61.69 | <0.0001 |
| DietNormal | -3.183 (0.558) | 73.55 | <0.0001 |
| BlockB | -0.556 (0.167) | 13.06 | <0.001 |
| InfectionInfected:Day2 | -0.077 (0.331) |  |  |
| InfectionInfected:Day3 | -0.309 (0.329) |  |  |
| InfectionInfected:Day5 | -0.067 (0.324) |  |  |
| InfectionInfected:Day7 | -0.593 (0.406) | 23.12 | <0.001 |
| InfectionInfected:DietNormal | 2.677 (0.638) | 24.99 | <0.0001 |
| Day2:DietNormal | 1.373 (0.580) |  |  |
| Day3:DietNormal | 0.509 (0.588) |  |  |
| Day5:DietNormal | 0.454 (0.591) |  |  |
| Day7:DietNormal | 1.289 (0.625) | 65.52 | <0.0001 |
| InfectionInfected:Day2:DietNormal | -0.717 (0.646) |  |  |
| InfectionInfected:Day3:DietNormal | -0.804 (0.659) |  |  |
| InfectionInfected:Day5:DietNormal | -1.792 (0.676) |  |  |
| InfectionInfected:Day7:DietNormal | -2.749 (0.777) | 22.37 | <0.001 |

